# A comparison of female competitive traits: Female aggression peaks at nest building but female song spans multiple contexts in a temperate songbird

**DOI:** 10.64898/2026.09.25.754226

**Authors:** Molly G. Gleason, Cara A. Krieg, Karan J. Odom

## Abstract

Female–female competition is increasingly recognized as a key driver of female ornamentation, including birdsong, which often functions in intrasexual competition. However, the specific resources females use elaborate traits to compete for remain unclear. In addition, few studies have simultaneously investigated the use of multiple competitive traits in females, despite growing independent interest in these traits (e.g., female song and aggression). We investigated the competitive contexts of female song, aggression and calling behavior in northern house wrens (*Troglodytes aedon*) to determine which resources females compete for across the breeding season. We simulated conspecific territorial intrusions using female song at three breeding stages representing different contexts: arrival (mate and territory acquisition), nest building (nest site and breeding status defense), and egg laying (brood defense). We tested whether female song and physical aggression varied as reproductive resources shifted across the breeding cycle. Females were significantly more aggressive during nest building, showing 5.8 times greater odds of a higher-intensity aggressive response during nest building compared to arrival. Female song output was similar across early stages but declined during egg laying, though this was not statistically significant after correction for multiple comparisons and individuals varied substantially in overall singing propensity. Non-song vocalizations varied by call type and breeding stage. Calls associated with aggression occurred most frequently during nest building, consistent with peak physical aggression responses. Together, these results identify nest building as the stage of highest female aggression, consistent with heightened competition over nest cavities and associated breeding status in this cavity-nesting species. In contrast, female song occurred across all stages and appears to function in multiple competitive contexts. This study provides evidence for context and mode-specific female signaling in a temperate songbird and highlights that females strategically use aggression, calls, and song to mediate social conflict across breeding contexts.

## Introduction

Traditional sexual selection theory has focused heavily on male traits and competition, often overlooking female roles in competitive mating contexts (Jones & Ratterman, 2009; Tobias et al., 2012). However, competition for resources like territories, mates, and food also drives the evolution of competitive or elaborate traits like female ornamentation and aggression (Andersson & Simmons, 2006; Clutton-Brock & Huchard, 2013; West-Eberhard, 1983). Female–female competition has direct fitness consequences, and traits such as song or aggression seem to evolve under sexual and social selection, just as in males. Females often compete for access to mates, high-quality territories, and limited nesting sites, particularly in species with ecological constraints or skewed parental investment (Clutton-Brock, 2009; Emlen & Oring, 1977; Tobias et al., 2012). Therefore, understanding when and how these traits are expressed is key to studying female signaling and competitive strategies (Cain & Rosvall, 2014). Thus, empirical studies evaluating the context and timing of female competition for status, resources, and reproductive access are necessary to determine the resources they are competing for (Rubenstein, 2012).

A key unresolved question is what specific resources females compete for across the breeding cycle. This competition is often mediated by elaborate, competitive traits like female birdsong and aggression, both of which are known to intensify during periods of heightened competition (Clutton-Brock, 2009; Langmore, 1998; Rosvall, 2011). Yet few studies distinguish whether these behaviors are driven primarily by a specific competitive context like mate acquisition or retention, nest defense, or brood protection, nor do they test how signaling shifts as the contested resource changes (Barros et al., 2024; Krieg & Getty, 2016; Rosvall, 2011). Additionally, most work evaluates female song or aggression in isolation and within a single context, making it difficult to determine how they function together across contexts. Because females face different selective pressures across the breeding cycle, understanding how they deploy vocal and non-vocal aggression across these contexts is essential to identify the specific resource that drives these competitive behaviors. These gaps highlight the need for explicit hypothesis testing that links signaling behavior to the underlying competitive stakes.

Moreover, when females use multiple signals, few studies have investigated whether these signals are used in independent or redundant ways. Here we evaluate how both female song and physical aggression function as competitive signals in different breeding contexts. Female song occurs most often in year-round territorial species and is associated with biparental care (Odom et al., 2025). Across species, it is used most often in territorial defense (Arcese et al., 1988; Barros et al., 2024), with females singing both individually (Illes & Yunes-Jimenez, 2008; Kriner & Schwabl, 1991) and in joint-defense with their mate (Logue & Gammon, 2004; Wheeldon et al., 2021). It is also clear that female song is often used in intrasexual competition (Barros et al., 2024; Krieg & Getty, 2016; Sierro et al., 2022). Females often respond more strongly to female song playback than to male song, showing sex-specific competitive signaling (Barros et al., 2024; Cain & Rosvall, 2014; Cooney & Cockburn, 1995; Krieg & Getty, 2016). Importantly, female songs in this context seem to have fitness benefits. In superb fairy-wrens (*Malurus cyaneus*), females that use song in intrasexual competition have higher reproductive success (Cain et al., 2015).

Females may also use song to compete for mates, sex-specific resources like a nest site, high-quality breeding territories, or to protect offspring (Austin et al., 2021; Krieg & Getty, 2016; Rosvall, 2011). In facultative polygynous species with biparental care, females often suffer a reduction in paternal care if their mate acquires a secondary female (Johnson et al., 1994; Llambias et al., 2012; Slagsvold & Lifjeld, 1994). In this case, females may use song to mate guard or defend access to paternal care (Langmore et al., 2002). However, female competition is also driven by nesting resources, with secondary cavity nesting species relying on limited nesting sites. Such species are particularly likely to engage in female competition far more than their open-cup nesting counterparts (Harris & Siefferman, 2014; Lipshutz & Rosvall, 2021). So, while it is clear that females compete, more research is needed to fully understand under which competitive contexts females invoke song.

Studies focused specifically on physical aggression in females offer further insight into the underlying competitive stakes. Also, like female bird song, female aggression across taxa has been historically understudied in relation to male aggression (Cain & Rosvall, 2014; Clutton-Brock & Huchard, 2013; Rosvall, 2011). Though males are often assumed to be the more aggressive sex, recent evidence shows that females can be as or more aggressive than males, especially when resources are limited, such as in cavity nesting species (Yu et al., 2025). Nonvocal aggressive behavior can include chasing, threat flights, physical attacks, and dive bombing and appears to serve different purposes across contexts (Rätti et al., 1994). It is shaped by both ecological and social pressures, with resource defense as a central theme across many species and systems (Stockley & Bro-Jørgensen, 2011; Tobias et al., 2012). In European starlings, more aggressive females were more successful in preventing their mates from becoming polygynous, maintaining monogamy through deterrence (Sandell, 1998). This pattern is not unique to birds: in prairie voles, female–female aggression rises specifically after a female pairs with a male, functioning as a form of mate guarding (Bowler et al., 2002). Female–female aggression in birds often peaks during periods of high reproductive investment and is especially intense during nest building, egg laying, or mate guarding (Sandell, 1998; Sandell & Smith, 1997). This link between aggression and reproductive investment is also seen in Drosophila, where females become markedly more aggressive after mating, as well as across species, where female aggressiveness scales with reproductive potential such as ovariole number (Bath et al., 2017; Bath & Gleason, 2025). However, further research is needed to explicitly test which of these resources drives aggressive behaviors, particularly in species where females face high competition across the breeding cycle.

Northern house wrens (*Troglodytes aedon*) are small, monomorphic, cavity nesting songbirds that breed across North America and are often socially monogamous but facultatively polygynous (Czapka & Johnson, 2000; Johnson et al., 1993). They provide an ideal system to explore the competitive contexts of female song and aggression because females face multiple limiting resources over the breeding season, including securing a mate, acquiring and defending nest cavities, and avoiding nest usurpation or ovicide by conspecifics (Johnson & Searcy, 1993; Krieg & Getty, 2020). Females reliably sing during the early breeding cycle, respond aggressively to female intruders, and respond more strongly to female song playback than to male playback, indicating that song functions in intrasexual competition (Gleason et al., *in press*; Johnson & Kermott, 1990; Krieg & Getty, 2016, 2020).

Although female house wrens likely use song and aggression to compete for mates, defend nesting cavities, or protect eggs and offspring against conspecific threats, these specific contexts have not been experimentally tested. We address this gap by conducting simulated territorial intrusions with female song playback across three breeding stages: arrival, nest building, and egg laying. House wrens exhibit a predictable breeding sequence that provides a natural experimental framework (Drilling & Thompson, 1988; Johnson & Kermott, 1991; Kendeigh, 1941). Each stage is marked by the resources a female stands to lose to an intruder. At arrival, when females select a male and his established territory, we hypothesize that competition at this stage centers on mate acquisition or retention. During nest building, once females have invested in lining a cavity, that cavity and lined nest are now at stake, plus the female’s breeding status on the territory. In this facultatively polygynous species, prospecting females can usurp other females’ nests, which results in not only loss of the nest when the female is physiologically ready to breed, but can also change her breeding status with the territorial male, who usually provides more care to the oldest nestlings on the territory (Dubois et al., 2006; Eckerle & Thompson, 2006; Johnson, 2024; Johnson et al., 1993). During egg laying, a female also stands to lose her eggs during a time when nests become vulnerable to ovicide plus usurpation by prospecting conspecific females (Belles-Isles & Picman, 1986; Freed, 1987). We hypothesize that the stage at which physical aggression and female song peak reflects when these combined resources are most worth defending.

In this study, we investigate the use of multiple competitive female behaviors in house wrens: songs, calls and physical aggression in response to a simulated female intruder across three breeding stages. In addition to asking when females are most responsive, we examine these three competitive behaviors to assess whether these behaviors act as a single integrated response or if they are deployed individually to defend different resources. We use differences among breeding stages and the resources at stake as a proxy for what females use each of these signals to compete for. If females use these behaviors primarily to *obtain* a mate and territory, we hypothesize these behaviors should peak during arrival. If nest cavities, a lined nest and breeding status are the resources most strongly defended, aggression and vocalizations should peak during nest building. Alternatively, if females use these competitive behaviors most to defend future offspring, responses should increase after egg laying. By identifying the stage at which females use these competitive behaviors most, we can infer which resources and breeding stages these behaviors function to defend. Overall, our study contributes to our understanding of elaborate, competitive female traits by identifying the specific resources that may select for these traits, as well as how multiple competitive behaviors may function in an individual female.

## Materials and Methods

### Study sites

We conducted simulated territorial intrusions at two sites in San Joaquin County, California: Heritage Oak Winery (HO; 38.155, -121.240) and Waldo Holt Preserve (WH; 38.237, -121.293). Heritage Oak Winery is a privately owned vineyard property bordered by riparian woodland along the Mokelumne River. Waldo Holt Preserve is reclaimed agricultural land containing mixed riparian forest, grassland, and seasonal wetland habitat located between Dry Creek and the Lower Sacramento River. At both properties, nest boxes were evenly spaced 30 m apart in transects along the tree line with 22 nest boxes at HO and 45 nest boxes at WH, for a total of 67 nest boxes. Both sites support high densities of breeding house wrens.

### Experimental design

We evaluated female house wren responses to simulated female house wren intrusions at breeding stages that represent key resources that female house wrens might compete for. These simulated intrusions were conducted at three stages of the breeding cycle, defined by observable transitions in female reproductive activity. We classified these experimental periods as (1) female arrival, (2) nest building, and (3) egg laying. We conducted arrival trials on the day of or within one day after a female was first observed at a territory, but before she began lining the nest (approximately 21 days before egg laying); nest building trials during the early nest-lining period, once females had begun adding soft material to the nest cup (approximately 11 days before egg laying), and egg laying trials during the early laying period, when 1–4 eggs were present in the nest.

Simulated intrusions consisted of a 3D-printed northern house wren model mounted on a pole with a high-fidelity INSMY C12 waterproof portable Bluetooth speaker hung immediately below the mount. We placed the mount and speaker within 1 m of the nest box. We created a 3D-printed house wren model for use in simulated intrusions to provide a target for any physical aggressive responses to female song playback. To create the model, we digitally scanned a taxidermy museum specimen (CAS ORN 92321) mounted in a typical wren posture with the tail cocked upward and facing forward (bill closed) using the Polycam 3D Scans app. The scan was processed in Blender to generate a printable model and base. The model was printed true to size (approximately 12 × 5 cm), based on measurements of the museum specimen and hand-painted to resemble a house wren (Figure S3). We used the same model across all trials to control for variation due to painting and to ensure that female song playback was the main signal tested in our playback experiments. We validated that house wrens responded to the model in preliminary observations before we began our simulated intrusions for the season. Responses were consistent with those described in prior studies using mounts. During experimental trials, a 10 m rope marked at 1 m intervals was laid on the ground, centered at the model, to estimate distances of up to 5 m in either direction of the model as focal birds approached. We calibrated playback volume to ∼85 dB SPL at 1 m using a digital sound-level meter, ensuring consistent broadcast amplitude across trials.

### Simulated territorial intrusion protocol

Each trial followed a modified protocol developed by Krieg and colleagues designed specifically for female northern house wrens to assess behavioral and vocal response to simulated territorial intrusions (Krieg & Burnett, 2017; Krieg & Getty, 2016). Following this protocol, trials began with approximately 5 minutes of setup, followed by 5–30 minutes of silence to confirm the presence of the focal female, a 5-minute playback period, and a 5-minute post-playback silence. If the focal female was not seen during the 5-minute set up period or for up to 30 minutes following set up, trials were not conducted. The observer was positioned 10–20 m from the nest in a portable hunting blind. All trials were recorded using a Tascam Portacapture X8 Linear PCM recorder paired with a Wildtronics omnidirectional microphone and parabolic reflector. Vocalizations were recorded continuously throughout the pre-playback, playback, and post-playback periods. The identity of responding birds was determined using unique color-band combinations and sex-specific vocalizations. The observer narrated the movements, distance to the model, and behaviors of all birds present. All simulated intrusions were conducted by a single observer to minimize inter-observer variation.

### Playback track preparation

We developed twenty-one unique playback tracks from focal female recordings collected in 2024 (the previous year) from both study sites. To reduce the possibility that the wrens in the current study were familiar with the wrens in the recordings, females were never assigned tracks from their own or neighboring territories. In addition, female house wrens in our population return to the same breeding site at low rates, so it was unlikely that a given female encountered her own or known songs in a subsequent breeding season. We derived 21 tracks from 11 individual females. For each focal female, we prepared two exemplars (“Track A” and “Track B”), each consisting of six unique songs produced by the same individual (except for one female, for whom only a single set of exemplar songs was available). We extracted individual songs in Raven Pro 1.6.5. To avoid pseudoreplication and ensure a balanced design, we rotated female tracks across trials and individuals so that exemplars were presented across experimental stages and no individual female heard the same playback more than once (Kroodsma, 1989; Kroodsma et al., 2001; McGregor et al., 1992). This helped avoid habituation of females in the study to the playbacks and ensured that no single playback disproportionately influenced results. Songs chosen for playbacks were approximately the same length (1.56 seconds with a standard deviation of 0.42 seconds). Each song was high-pass filtered at 1.5 kHz to remove low-frequency background noise and normalized to –1 dB to standardize playback amplitude. Final tracks consisted of six songs separated by 5-second silent intervals, looped continuously for 5 minutes. We edited all playback tracks in Adobe Audition (Adobe Inc., 2025).

### Sample sizes

We conducted simulated intrusions across three experimental stages: arrival (n = 15), nest building (n = 15), and egg laying (n = 20). In total, we tested 29 unique females across 50 trials (Table 1). Of these, 12 trials were from Heritage Oak Winery (HO) and 38 trials were from Waldo Holt Preserve (WH). Because our experimental design was based on the breeding stage, naturally, all females tested at the arrival stage were naïve to simulated intrusions. To control for potential order effects across breeding stages, we conducted a subset of trials at the nest building and egg laying stages on naïve females that had not previously experienced simulated intrusions. In the nest building stage, 9 females were naïve and 6 had previously received arrival trials. In the egg laying stage, the sample comprised 5 naïve females and 15 that previously received playback trials. Of these, 5 had received nest building only, 4 had received arrival trials only, and 6 had received both previous stages (arrival and nest building). No female ever received the same playback exemplar (Track A or Track B from a given individual) at multiple stages.

**Table 1.** Number of simulated intrusion trials by site (Heritage Oak Winery, HO; Waldo Holt Preserve, WH) and experimental stage (arrival, nest building, egg laying).

| Stage | HO Trials | WH Trials | Total Trials |
| --- | --- | --- | --- |
| Arrival | 3 | 12 | 15 |
| Nest building | 4 | 11 | 15 |
| Egg laying | 5 | 15 | 20 |
| Total | 12 | 38 | 50 |

### Nonvocal response to simulated intrusions

We quantified female nonvocal response to simulated intrusions using three behavioral metrics adjusted from Krieg & Getty (2018, 2020): (1) **Dives or attacks**. Dives were defined as rapid swoops passing within 1 m of the speaker/model. Attacks were considered any physical contact like pecking or landing on the model; (2) **Latency to first approach within 5 m** of the model measured from the start of the playback. If a female began inside 5 m, latency was the time to move closer or to re-enter the 5 m radius after any initial retreat. Because the combined playback and post-playback period lasted 600 s, females that never entered the 5 m radius during that time were assigned a latency of 600 s; and (3) **Presence within 1 m of the model** during the playback or post-playback period when not already within 1 m during the pre-playback period, recorded as a yes/no metric. Approaching this closely places the female at heightened risk of physical attack if the model were a real rival. This controlled for birds that perched near the nest box in non-agonistic contexts and distinguishes those that retreated when the playback started.

These behaviors are commonly used in playback studies as reliable indicators of aggression, particularly in same-sex competition where mating motivations are not driving the response. To capture the full range of behavioral responses and enable comparison across individuals and breeding stages, we combined these measures into an ordinal aggression index. Females were classified into three categories of escalating aggression (Table 2).

**Table 2.** Aggression classification of female response to simulated intrusions.

| Aggression Class | Criteria |
| --- | --- |
| Low | Bird never dove at or attacked the model.<br>Bird did not approach within 5 m during the entire playback period.<br>Bird spent no time within 1 m of the model during the playback period. |
| Medium | Bird never dove at or attacked the model.<br>Bird spent no time within 1 m of the model.<br>Bird approached within 5 m during the playback period but took > 30 s. |
| High | Met any of the following:<br>1) Bird attacked or dove at the model<br>2) Bird approached within 5 m with a latency $\leq 30$ s<br>3) Bird entered within 1 m when not already within 1 m during the pre-playback period. |

To verify that our Low-Medium-High aggression classes reflected a single underlying aggression gradient, we conducted a principal component analysis (PCA) on the three behavioral measures. PC1 scores increased monotonically across the three classes (Table S1), confirming that the categories captured a common aggression axis.

### Quantification of vocal response to simulated intrusions

Northern house wrens produce sex-specific song (Johnson & Kermott, 1990; Keck et al., 2025; Krieg & Getty, 2016). House wrens also produce a variety of broadband calls (e.g., chatters, chutters, churrs, rattles) whose structure and context remain incompletely described, alongside sex-specific calls such as male squeaks and female distress or whine calls (Johnson, 2024). Vocalizations were assigned to eight recognizable types based on distinct structural differences observed in spectrograms, encompassing the full range of acoustic variation recorded during simulated intrusions (Table 3; Figure 1). Several call types have established functions in house wrens or closely related contexts: HI calls are associated with aggressive interactions in females, occurring more frequently during conspecific playback and predicting subsequent physical attacks (Krieg & Burnett, 2017), while harsh calls and rattles have been documented in predator response and territorial defense respectively (Johnson, 2024). Sweep calls, though structurally similar to high-pitched squeaks described in males during courtship (Johnson, 2024), were observed here during aggressive interactions by females and have not been previously described in females. The functional context of mew and quack calls remains undescribed. Full acoustic descriptors, frequency ranges, and quantification methods for all vocalizations can be found in Table 3.

**Figure 1.**
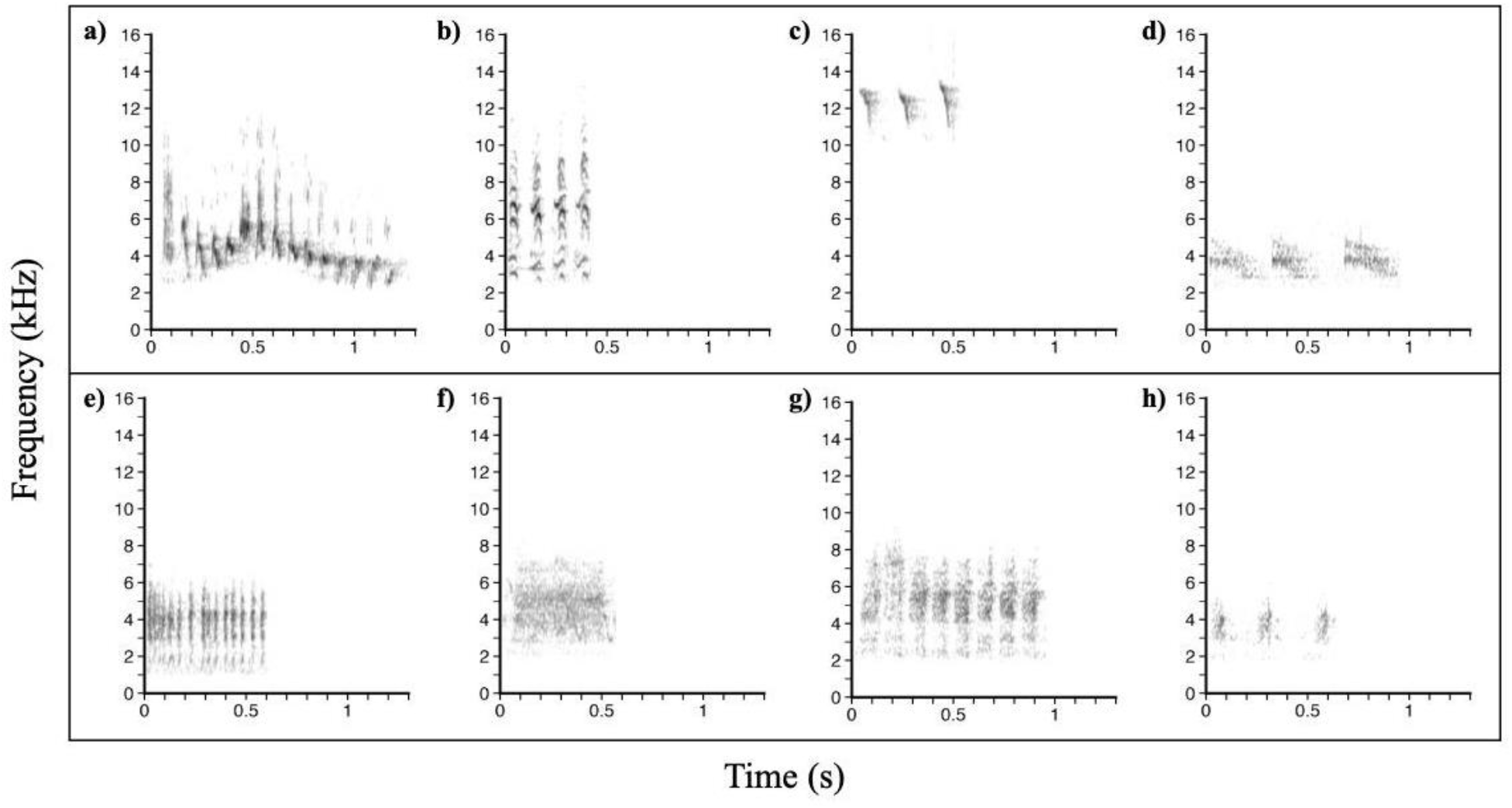
Representative spectrograms of female house wren vocalizations. (a) female song (b) four individual HI calls (c) three sweep calls (d) three individual harsh calls (e) chatter burst (f) one mew call (g) eight rattle notes (h) three quack calls. Darkness is not indicative of absolute differences in amplitude across vocalizations.

**Table 3.** Classification of female house wren vocalizations.

| Vocalization Type | Frequency range (kHz) | Acoustic features | Quantification unit | Known context |
| --- | --- | --- | --- | --- |
| Female Song | 2–11 | ~1 s, broadband with harmonics; variable complexity | Song unit | Territorial and competitive contexts (Johnson & Kermott, 1990; Krieg & Getty, 2016) |
| HI call | 3–14 | Very short (~0.04 s), low-amplitude, high-frequency notes with multiple harmonic overtones; produced singly or in short series | Individual notes | Aggressive interactions; predicts physical attack (Krieg & Burnett, 2017); interactions between mates during courtship and incubation (Krieg, pers. obs.) |
| Sweep | 9–14 | Very short (~0.08 s), high-frequency, tonal call that sweeps downward in frequency | Individual notes | Similar to male courtship squeaks (Johnson, 2024); observed in aggressive contexts (pers. obs.) |
| Harsh | 3–6 | Short (~0.3 s) broadband, noisy notes | Individual notes | Predator response near nest, both sexes (Johnson, 2024) |
| Chatter | 1–7 | Bursts of very short (~0.03 s), very closely spaced notes | Duration (s) of bouts ($\leq 50$ ms apart) | Broadly described for species; context incompletely defined, likely involved in predator response near nest (Johnson, 2024) |
| Mew | 2–7 | Simple hissy/buzzy call (~0.5 s), catbird-like quality | Individual notes | Context unknown |
| Rattle | 2–8 | Mechanical trill of closely spaced notes (~0.05 s apart) | Individual notes | Territorial defense and aggression (Johnson, 2024) |
| Quack | 2–6 | (~0.075 s) nasal, duck-like calls, variable repetition | Individual notes | Context unknown |

All vocalizations were manually annotated in Raven Pro (v. 1.6.5; Cornell Lab of Ornithology) using selection tables to document the call type and start and end time for each vocalization. These annotations were then exported and compiled in R Studio (v. 4.6.0; R Core Team, 2026) into a dataset summarizing call tallies by trial and playback stage. The final dataset included 50 trials from 29 unique females, each with quantified totals of all eight vocalization types per experimental stage.

### Analysis

We conducted three separate analyses to assess female responses to simulated rivals across breeding stages: (1) non-vocal aggression, (2) female song, and (3) other vocal responses. We performed all analyses in R (R Core Team, 2026). Across analyses, the breeding stage was the primary predictor, and we used mixed-effects models with Female ID as a random effect to account for repeated measures. Aggression was tested using ordinal aggression scores, female song was analyzed as the number of songs produced, and principal components analysis was used to summarize variation in call types. To address the modest sample size, we examined effect sizes (odds ratios for aggression and back-transformed rate ratios with 95% intervals for song) alongside significance.

We evaluated *Site* and *Playback Track* as potential covariates in our models. *Site* was retained only in the final model for female song output because it improved model performance, whereas *Playback Track* did not improve fit for any response and was excluded from final models. Model selection was based on information-theoretic or cross-validation criteria (AIC or PSIS-LOO), and model assumptions were assessed through residual diagnostics. The Bayesian aggression model used 95% credible intervals and the probability of direction, with a derived two-sided p-value. We treated α = 0.05 as the threshold for statistical significance. For female song output and call PC scores, all pairwise post-hoc contrasts were adjusted for multiple comparisons using Tukey’s HSD. Pairwise post hoc contrasts were obtained using the *emmeans* package (Lenth, 2025). The female song figure reports observed counts, whereas aggression and call figures report estimated marginal means ± 95% credible or confidence intervals.

To assess whether responses changed with repeated simulated territorial intrusion trials, we tested the effect of within-individual playback trial number (1–3) in separate preliminary models. Trial number did not significantly predict any behavioral or vocal response and was therefore excluded from the final models.

### Physical Aggression analysis

To test whether female physical aggression levels varied across breeding stages, we analyzed the aggression categories using a Bayesian cumulative-logit mixed model which allowed us to handle an ordinal variable in a mixed model format. Aggression was treated as an ordered response (*Low*, *Medium*, *High*) with *Experimental Stage* as a fixed effect and a random intercept for *FemaleID* to account for repeated measures. Models were fit using the *brms* package (Bürkner, 2017), which interfaces with Stan (Guo et al., 2025; Stan Development Team, 2020). We used weakly informative Normal(0, 2.5) priors on fixed-effect coefficients and brms default Student-t(3, 0, 2.5) priors on thresholds and the random-effect standard deviation. Four Markov-chain Monte Carlo (MCMC) chains were run for 2,000 iterations each (1,000 warm-up). Convergence was assessed using R-hat values (< 1.01; Vehtari et al., 2021) and effective sample size diagnostics. Pairwise contrasts between breeding stages were computed from the posterior distribution using the emmeans package (Lenth, 2025). For each contrast, we reported the posterior median, 95% equal-tailed credible interval (CrI), the posterior probability that the effect was in the estimated direction [P(direction)], and a derived two-sided p value calculated as 2 x (1-P(direction)). We used equal-tailed intervals so that the reported CrI, pd, and p-value are consistent. To verify our results, we repeated the analysis using the continuous aggression score (the first principal component of the three aggression metrics; Table S1). This analysis produced the same pattern as the Bayesian cumulative-logit mixed model (Table S4).

### Female song analysis

To test whether female song output differed across breeding stages, we modeled the total number of songs produced during playback and post-playback periods using a generalized linear mixed model in *glmmTMB* (Brooks et al., 2017) with a Tweedie distribution to handle overdispersed, zero-heavy counts. Model fit was assessed with the *DHARMa* package (Hartig et al., 2024) by simulating 1,000 residual datasets through *simulateResiduals*(), which showed no major deviations from model assumptions. *Site* was retained in the final model because it improved model fit (lowest AIC). To quantify the magnitude of stage differences in female song output, we calculated model-predicted effect sizes using estimated marginal means derived from the fitted Tweedie generalized linear mixed model. Pairwise contrasts were back-transformed to the response scale to obtain ratio differences in predicted song counts among breeding stages, along with 95% confidence intervals.

### Other vocal response analysis

To evaluate differences in non-song vocalizations (HI, Sweep, Harsh, Chatter, Mew, Quack, Rattle) across breeding stages, we first pooled rare call types (Mew, Quack, Rattle) into a single “Other” category. Call counts and durations were logₑ(x + 1)–transformed using *log1p()* to reduce skew and stabilize variance. We then conducted a principal components analysis (PCA) on transformed totals using the *principal()* function in the *psych* package (Revelle, 2026). Because the extracted components were correlated (|r| up to 0.35), we applied an oblique (promax) rotation (Hendrickson & White, 1964) and retained three components. The rotated components explained ∼83% of variance: PC1 (HI, Sweeps, Harsh), PC2 (Chatter, Harsh), and PC3 (Rare ‘Other’ calls) (Table S7, S8). Differences in component scores across breeding stages were then tested using linear mixed models.

### Behavioral correlations

Finally, we examined associations between aggression class and vocal behavior by testing relationships between the ordinal aggression categories (coded 1–3, from low to high) and each vocalization type (female song, HI, Sweeps, Harsh, Chatter, Mew, Rattle, Quack). We tested correlations between aggression scores and the rate of each vocalization using Spearman rank correlation. We corrected for multiple comparisons for these analyses using the Bonferroni method.

### Ethical note

This research was approved by the (Redacted) Institutional Animal Care and Use Committee (IACUC approval number 00000016) and followed the Guidelines for the Use of Wild Birds in Research established by the Ornithological Council. Birds were banded under Federal Bird Banding Permit number 22712 (under Master Bander T. Coombes-Hahn). Birds were either hand-trapped on the nest during incubation or in mist nets that were monitored continuously. All individuals were processed (measured, banded and blood drawn) and released at the territory of capture as quickly as possible. Simulated territorial intrusions were limited to 15 minutes per nest box per day to minimize disturbance and we avoided conducting consecutive trials at neighboring nest boxes to minimize disturbance and potential compounding effects. No birds abandoned territories following processing or behavioral trials. No birds suffered adverse effects during brief capture and measurement.

## Results

In response to the simulated female rival trials, females showed the strongest physical aggressive responses during the nest building stage (Figure 2; Table 4). Females tended to sing more during arrival and nest building than during egg laying, but neither difference was statistically significant after correction for multiple comparisons (Figure 3; Table 6). Overall vocal output (both songs and calls) declined by egg laying, with other call types reaching their lowest in the final stage (Figure 4; Table 7). Among vocalizations, HI calls and Sweep calls were the strongest correlates with female aggression, showing moderately to strongly positive associations with aggression scores (Figure S2). These aggressive calls also occurred most during the nest building stage (Table 7).

**Figure 2.**
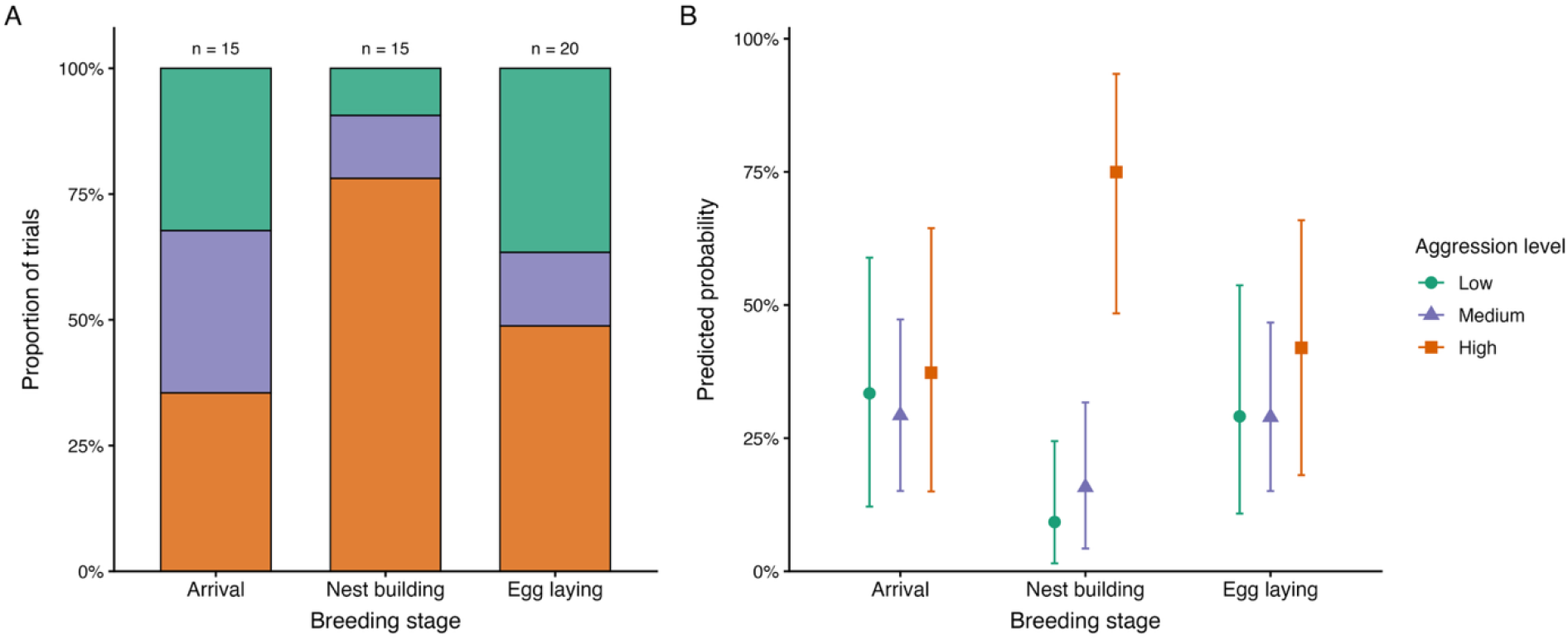
Female aggressive response to simulated intrusions across breeding stages. A. Observed proportions of physical aggression classes (low, medium, high) during trials across breeding stages. B. Predicted posterior probabilities of aggression classes across breeding stages derived from a Bayesian ordinal mixed-effects model.

**Figure 3.**
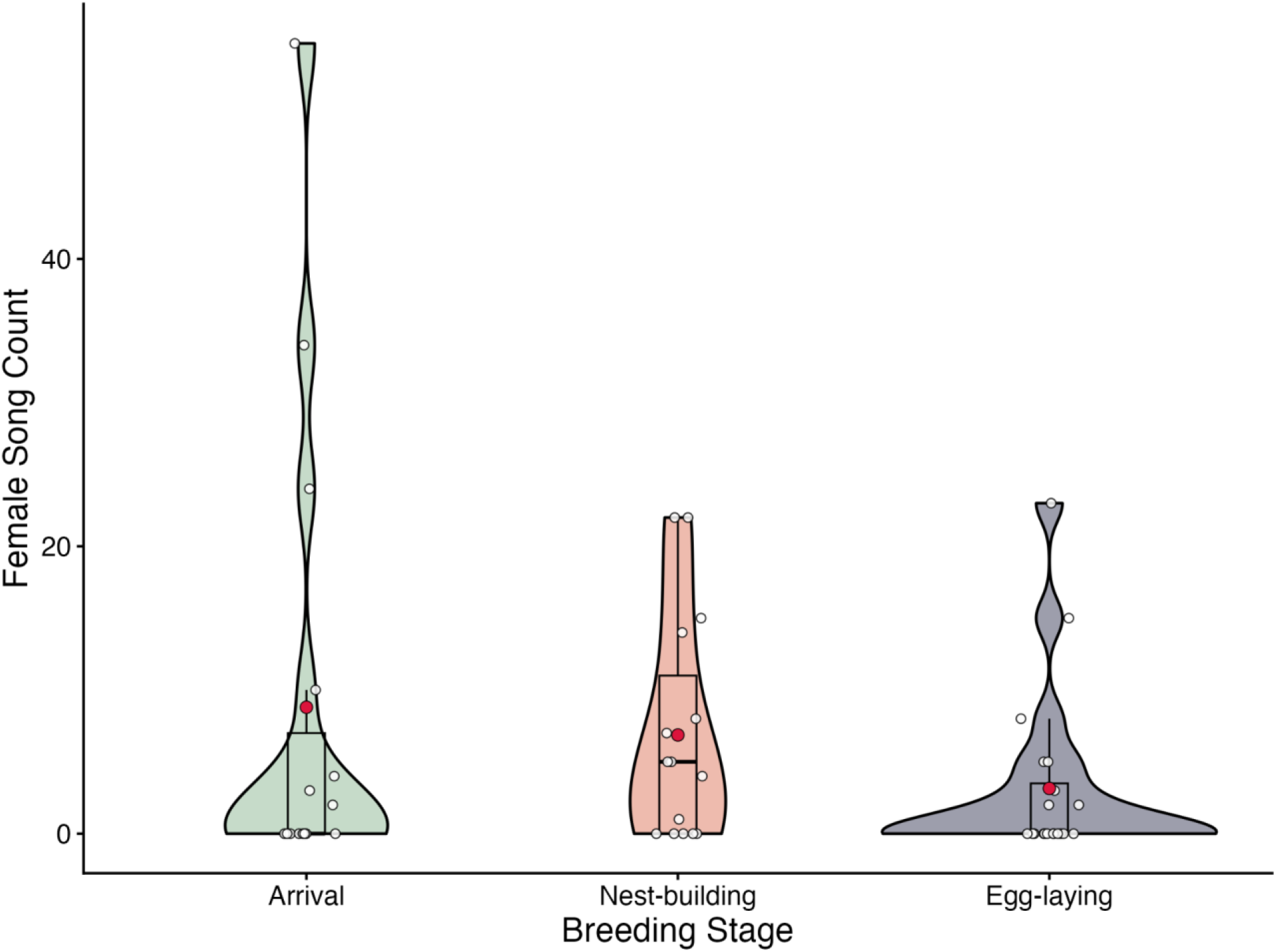
Female song responses to simulated intrusions across breeding stages. Violin plots show the distribution of song counts across breeding stages (arrival, nest building, and egg laying). White circles represent individual females; red circles indicate group means; and black bars denote interquartile ranges. Arrival n =15; Nest building n=15; egg laying n=20).

**Figure 4.**
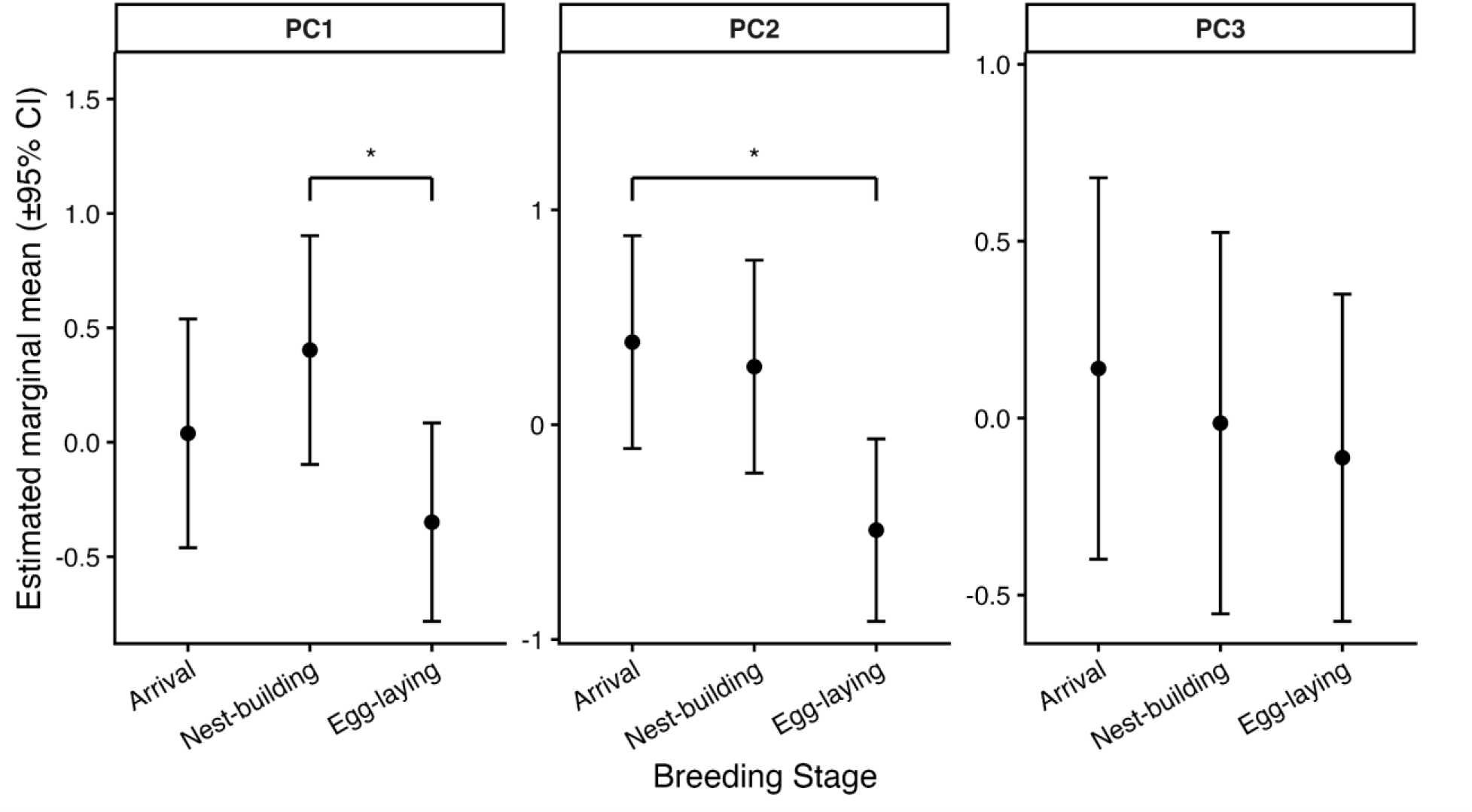
Call PC scores across breeding stages. Estimated marginal means (±95% CI) of PC1–3 across arrival, nest building, and egg laying stages, derived from linear mixed-effects models. PC1 loads primarily on calls associated with aggressive contexts; PC2 on scolding calls; PC3 on rare call types. Asterisks indicate significant pairwise differences based on Tukey-adjusted comparisons. Arrival n =15; Nest building n=15; egg laying n=20).

**Table 4.** Breeding-stage differences in female aggression intensity (Low → Medium → High) from a Bayesian cumulative-logit mixed model with female identity as a random intercept. Estimates are posterior medians (log-odds); SE is the posterior dispersion; 95% CrI is the equal-tailed credible interval; odds ratios are exp(estimate); the Bayesian probability of direction; and a frequentist-style p-value calculated as 2 × (1 − pd). Bold rows indicate p < 0.05. Based on 50 trials from 29 females.

| Variable | Estimate | SE | 95% CrI | Odds ratio | P(direction) | p-value |
| --- | --- | --- | --- | --- | --- | --- |
| Intercept (Low vs Medium) | −0.73 | 0.59 | [−1.98, 0.36] | — | — | — |
| Intercept (Medium vs High) | 0.57 | 0.60 | [−0.54, 1.79] | — | — | — |
| **Nest building vs Arrival** | **1.76** | **0.83** | **[0.24, 3.49]** | **5.81** | **0.988** | **0.025** |
| **Nest building vs Egg laying** | **1.51** | **0.84** | **[0.01, 3.37]** | **4.52** | **0.976** | **0.048** |
| Arrival vs Egg laying | −0.23 | 0.69 | [−1.56, 1.13] | 0.79 | 0.626 | 0.748 |

### Physical Aggression

Females showed the highest physical aggression levels during nest building (Figure 2).

Compared to arrival, females at nest building were significantly more aggressive (log-OR = 1.76, 95% CrI [0.24, 3.49]; P(direction) = 0.99; p = 0.025), corresponding to roughly 5.8 times greater odds of a higher-intensity aggressive response (Table 4). Aggression at nest building also exceeded that during egg laying (log-OR = 1.51, 95% CrI [0.01, 3.37]; P(direction) = 0.98; p = 0.048); corresponding to approximately 4.5 times greater odds. Arrival and egg laying did not differ in aggression level (log-OR = −0.20, 95% CrI [−1.56, 1.13]; P(direction) = 0.63; p = 0.748). The random effect of female identity (σ = 1.05, 95% CrI [0.07, 2.57]) indicated moderate among-female variation in baseline aggression.

**Table 5.**
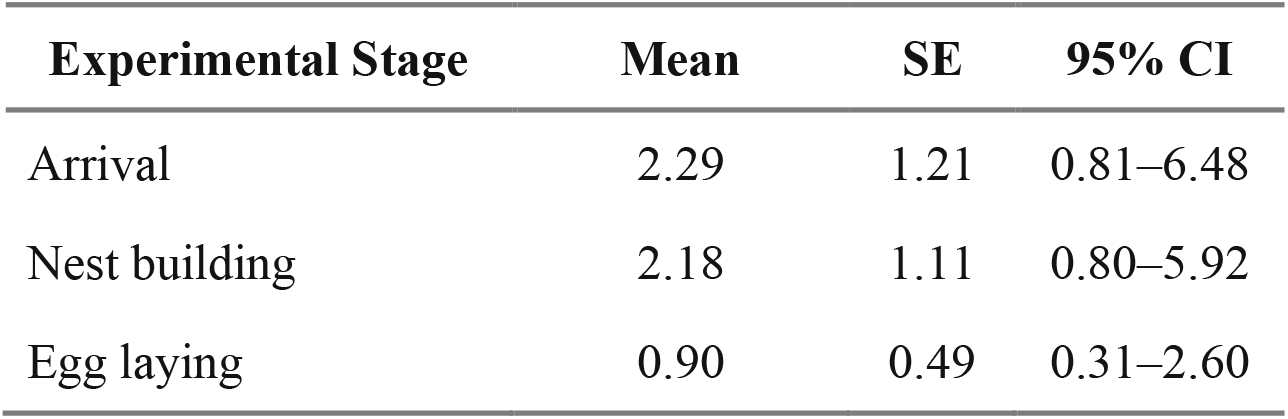
Model-estimated female song output during simulated intrusions across breeding stages. Values are estimated marginal means (back-transformed) from a Tweedie generalized linear mixed model (log link) with female identity as a random effect. SE and 95% CI reflect model-based uncertainty on the response scale.

### Song

Female song output was generally similar during the early breeding stages but declined by the egg laying period (Figure 3). Model-predicted female song output was lowest during egg laying, with females predicted to produce approximately 2.29 ± 1.21 songs during arrival, 2.18 ± 1.11 songs during nest building, and 0.90 ± 0.49 songs during egg laying across playback and post-playback periods (Table 5). Female song output tended to be lower during egg laying than during arrival. Compared to arrival, female song output during nest building was similar (log-scale estimate = −0.05 ± 0.42, z = −0.12, p = 0.90; Table S6). Song output during egg laying compared to arrival was initially significant in the uncorrected model (log-scale estimate = −0.93 ± 0.44, z = −2.13, p = 0.033; Table S6), however this difference was not significant after correcting for multiple comparisons (Tukey-adjusted p = 0.084; Table 6).

**Table 6.** Pairwise breeding stage contrasts for female song output from the Tweedie generalized linear mixed model. Estimates are on the log scale (first stage minus second); rate ratios are exp(estimate). 95% CIs and p-values are Tukey-adjusted across the three comparisons. No contrast reached p < 0.05. Full model coefficients are in Table S6.

| Contrast | Estimate | SE | 95% CI | Rate ratio | z-score | p-value |
| --- | --- | --- | --- | --- | --- | --- |
| Arrival vs Nest building | 0.05 | 0.42 | [−0.93, 1.04] | 1.05 | 0.12 | 0.992 |
| Arrival vs Egg laying | 0.93 | 0.44 | [−0.09, 1.96] | 2.54 | 2.13 | 0.084 |
| Nest building vs Egg laying | 0.88 | 0.42 | [−0.11, 1.87] | 2.41 | 2.09 | 0.092 |

However, contrasts on the response scale indicated that predicted song output during arrival and nest building was approximately 2.4–2.5 times higher compared to egg laying, whereas song output was similar between arrival and nest building (Table 6). Although these differences were not statistically significant, their consistent direction suggests a decline in female song output by the egg laying stage.

### Calls

Total vocal output and call type use varied across breeding stages (Figure 4). A Promax-rotated PCA extracted three components that together explained 82.5% of the total variance in vocal behavior (Table S7). PC1 (36.2%) described variation in vocalizations associated with aggressive contexts (sweeps, HI, harsh calls), PC2 (23.9%) captured broadband scolding vocalizations (chatter, harsh calls), and PC3 (22.3%) represented rare call types.

Females produced the greatest number of calls associated with PC1 during the nest building stage (Figure 4). PC1 captured calls that appear to be associated with aggressive contexts: HI, Sweeps, and Harsh calls. HI (0.82), Sweeps (0.93), and Harsh (0.54) all loaded strongly and positively, with higher PC1 scores indicating greater production of these call types (Table S7). Females at nest building had the highest PC1 scores, those at arrival were intermediate, and egg laying females had the lowest scores, indicating a decline in HI/Sweep/Harsh calling by the final stage (Type III ANOVA, F = 3.29, p = 0.049; Table 7). Tukey post hoc tests confirmed a significant decrease from nest building to egg laying (p = 0.046); other contrasts were not significant (Table 7).

**Table 7.** Pairwise breeding stage contrasts for call PC scores from linear mixed models with female identity as a random intercept. Each PC’s overall stage effect (Type III F-test, Satterthwaite df) is given in the section headers. Contrast estimates are differences in PC scores; 95% CIs and p-values are Tukey-adjusted across the three comparisons within each PC. Bold rows indicate p < 0.05.

| Contrast | Estimate | SE | 95% CI | t-value | p-value |
| --- | --- | --- | --- | --- | --- |
| *PC1 —overall stage effect: $F(2, 35.2) = 3.29, p = 0.049$* |  |  |  |  |  |
| Arrival vs Nest building | -0.36 | 0.33 | [-1.18, 0.45] | -1.09 | 0.525 |
| Arrival vs Egg laying | 0.39 | 0.30 | [-0.36, 1.13] | 1.28 | 0.417 |
| **Nest building vs Egg laying** | **0.75** | **0.30** | **[0.01, 1.49]** | **2.50** | **0.046** |
| *PC2 —overall stage effect: $F(2, 47) = 4.67, p = 0.014$* |  |  |  |  |  |
| Arrival vs Nest building | 0.11 | 0.35 | [-0.73, 0.96] | 0.33 | 0.942 |
| **Arrival vs Egg laying** | **0.88** | **0.32** | **[0.08, 1.67]** | **2.71** | **0.027** |
| Nest building vs Egg laying | 0.76 | 0.32 | [-0.03, 1.55] | 2.36 | 0.062 |
| *PC3 —overall stage effect: $F(2, 39.8) = 0.27, p = 0.764$* |  |  |  |  |  |
| Arrival vs Nest building | 0.15 | 0.38 | [-0.76, 1.07] | 0.41 | 0.911 |
| Arrival vs Egg laying | 0.25 | 0.35 | [-0.60, 1.10] | 0.73 | 0.751 |
| Nest building vs Egg laying | 0.10 | 0.35 | [-0.75, 0.95] | 0.28 | 0.957 |

**Table 8.**
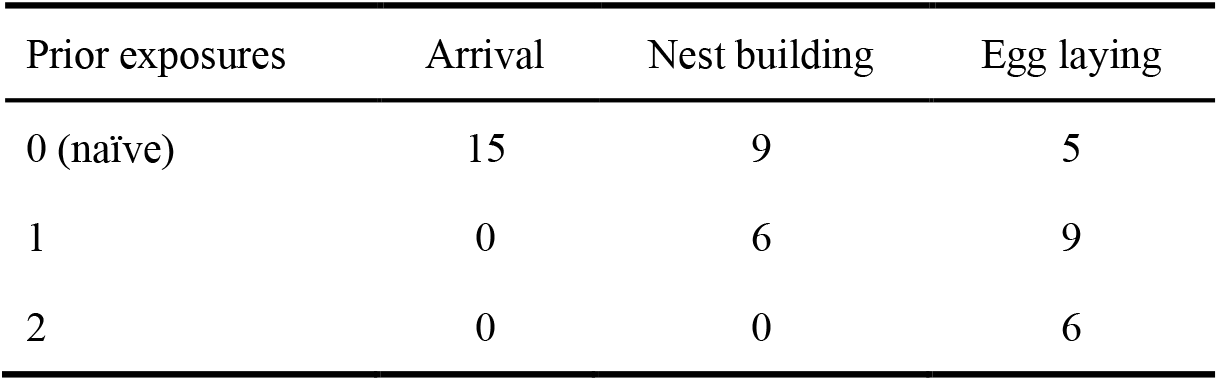
Number of females at each breeding stage by prior simulated intrusion exposure. Values indicate how many females had experienced zero, one, or two previous simulated intrusion trials at the time of testing.

| Prior exposures | Arrival | Nest building | Egg laying |
| --- | --- | --- | --- |
| 0 (naïve) | 15 | 9 | 5 |
| 1 | 0 | 6 | 9 |
| 2 | 0 | 0 | 6 |

Females produced more PC2 associated calls upon arrival and reached the lowest levels during egg laying. PC2 captured scolding calls, with chatter loading strongly (1.02) and harsh calls contributing modestly (0.41). Higher PC2 scores reflected greater production of these call types. PC2 varied among stages (Type III ANOVA, F = 4.67, p = 0.014; Table 7), and females at arrival had higher PC2 scores than those at egg laying (Tukey p = 0.027; Table 7). Other pairwise contrasts were not significant.

Rare call types did not differ across breeding stages. PC3 included ‘Other’ call types, loading very strongly (0.99) and a smaller loading of Sweep calls (0.30). PC3 scores did not differ across breeding stages (Type III ANOVA, F = 0.271, p = 0.764).

### Relationship between aggression and female vocalizations

To investigate how female vocal behavior was associated with aggressive behavior, we tested correlations between aggression scores and the rate of each vocalization type. Sweep calls and HI calls had the strongest association with female aggression, with sweep calls moderately to strongly correlated with aggression (ρ = 0.53, p < 0.0001) and HI calls moderately correlated (ρ = 0.46, p = 0.0009; Figure S2). Sweep (ρ = 0.53) and HI (ρ = 0.46) remained significant after multiple-comparison correction (adjusted p ≤ 0.008). Female song showed a weak, non-significant trend toward higher aggression (ρ = 0.27, p = 0.062). No meaningful relationships were detected for harsh, chatter, mew, rattle or quack calls.

### Effect of repeated simulated intrusion trials within individuals

To assess whether repeated exposure to simulated intrusions affected female responses, we tested whether trial number (i.e., how many prior trials a female had experienced) predicted song output, aggression, or call output. Responses did not change systematically with trial number: song output (Tweedie GLMM: β = –0.39 ± 0.27 SE, p = 0.15), aggression intensity (β = –0.07, 95% CrI [–0.96, 0.83]), PC1 (β = –0.28 ± 0.18 SE, p = 0.14), and PC3 (β = 0.08 ± 0.20 SE, p = 0.71) were all unaffected. PC2 showed a negative relationship with trial number in an initial model (β = –0.46, p = 0.022), but this disappeared once breeding stage was included (β = – 0.17, p = 0.486), suggesting the pattern reflected stage-related variation rather than habituation or sensitization.

### Response rates

Most of our experimental trials elicited a behavioral response. Nearly all females approached the simulated intruder during nest building (14 of 15 trials elicited an approach within 5 m), compared with 10 of 15 at arrival and 13 of 20 at egg laying. Many non-approaching females still responded vocally, and only 8 of 50 trials (16%) elicited no response on any measure. For all playback trials, we confirmed that the female was present on the territory the morning of or shortly before beginning the experimental trial. Therefore, we considered a lack of response as a genuine low response and kept these data. The lower approach rates at arrival and egg laying are consistent with the stage-dependent aggression findings.

## Discussion

Our results revealed that female house wrens respond differently to conspecific female intruders depending on the breeding stage, and that they use their separate competitive signals differently in nuanced ways. Notably, females were most physically aggressive during nest building compared to arrival and egg laying. Similarly, females produced the highest number of calls known to be used in aggressive contexts during the nest building stage. Interestingly, female song output in response to territorial intrusions did not differ significantly between breeding stages, though there was a nonsignificant trend that some females tended to sing more upon arrival. In addition, we found that vocal output was lowest during the egg laying stage. Overall, this study helps us understand how females use competitive behaviors across breeding stages and identify which resources most strongly drive specific aspects of female-female competition.

### High female aggression at nest building

Our results showed that females were most aggressive during the nest building stage. In many systems, female competition arises because access to essential breeding resources is limited, selecting for behaviors that allow females to compete for these resources (Clutton-Brock & Huchard, 2013; Rosvall, 2011; Stockley & Campbell, 2013). We hypothesize that each breeding stage represents different resources that females compete for throughout the breeding cycle: (1) mate attraction or competition and securing a territory, (2) established nest defense and maintaining their breeding status, and (3) brood protection. We therefore predict that female-female competitive behavior in house wrens will peak at the stage where the most highly defended resources are at stake. Because aggression peaked at nest building, we interpret our findings to mean that access to the nest cavity a female has already lined and invested in and maintaining breeding status are the main focus of competition at this stage. If females are mainly competing for a nest site and not a mate or defending offspring, females should be most aggressive when the risk of losing the cavity is the highest. During female arrival, territories are not fully established. Many females have still yet to arrive at the breeding grounds; most males are still unmated, and many nest boxes are still unclaimed. Females may also take several days to decide on a territory/male. It is not uncommon to spot the female on multiple territories soon after arrival before breeding is more imminent and she is consistently spotted in one location (Krieg, pers. obs.). If a female lost the nest site at this stage, she would be able to obtain a new nest site relatively easily. By nest building, however, females have invested both time and resources into the nest site. In addition, many cavities in preferred habitat have been taken while others are still being competed for. This makes established nest sites vulnerable to takeover by females who are still actively searching for nesting sites. If a female loses her cavity at this stage, she may be unable to secure another in time to breed within that reproductive cycle, when she may be within days of being reproductively ready to lay eggs. This mirrors female jumping spiders, where high valuation of contested resources, rather than fighting ability, drives the escalated ’desperado’ tactics expected when the stakes of losing are high (Elias et al., 2010).

Additionally, the threat an intruding female poses to the resident female’s breeding status may also be greatest at this stage. In populations where facultative polygyny is common, an additional prospective female is most threatening during nest building, as the two females’ breeding cycles have potential to overlap most closely. This can intensify competition for male attention as the primary versus secondary female status is not yet resolved. By egg laying, even if a male acquired a second female, the resident female has secured her position as the primary female. Being the first female to breed on a territory offers significant advantages as male house wrens provide more provisioning to the oldest nest on the territory (Czapka & Johnson, 2000; Johnson et al., 1993). Defense of breeding status would therefore predict higher aggression during nest building and lower aggression during egg laying. However, polygyny appears to be low or absent in our California study population, in contrast to the higher rates reported in some eastern and midwestern populations (Gleason et al., *in press*). The reduced polygyny in our population suggests breeding-status defense may not be a main driver and instead further suggests that the nest building peak in aggression reflects competition for continued cavity and nest access.

This pattern is especially pronounced in cavity-nesting species, where suitable nesting sites are often limited and aggressively defended. In many species across taxa, female aggression is essential for securing a nesting territory (Albers et al., 2017; Duckworth & Badyaev, 2007; Lipshutz & Rosvall, 2021; Sandell & Smith, 1997; Schuppe et al., 2016; While et al., 2009; Wu et al., 2019). Cavities are actively contested and stolen across cavity nesting species (Cornelius et al., 2008; Wiebe, 2011). For example, nest usurpation is common in tree swallows, where aggressive interactions help individuals retain access to nest cavities (Leffelaar & Robertson, 1985; Rosvall, 2008). In general, cavity nesting birds show greater conspecific territorial aggression than closely related species with alternative nesting strategies (Lipshutz & Rosvall, 2021), and this aggression is frequently directed toward female intruders (Yu et al., 2025). The same dynamic appears outside birds wherever breeding sites are scarce and reusable. In shell-dwelling cichlids, females compete over and take over the empty snail shells they need as brood chambers (Gübel et al., 2021; Walter & Trillmich, 1994). More aggressive female northern house wrens are also less likely to lose their entire clutch during territorial replacement by other wrens when competition for cavities is experimentally elevated (Krieg & Getty, 2020). Furthermore, additional work in northern house wrens suggests that females value nest sites more strongly than access to high-quality males or paternal care. When researchers experimentally added supplemental nest boxes, females with access to additional cavities showed significantly lower aggression toward simulated intruders compared to females in territories without additional boxes (Krieg & Getty, 2018), suggesting that nest cavities are the primary resource under competition. Our finding that female aggression peaked during nest building, when cavities in preferred habitat are most actively contested, further supports this conclusion. An important caveat is that breeding stage serves as a proxy for the resource at stake. Because we did not measure individual variation in territory, nest, or mate quality, we cannot fully separate what females compete for from how much a given female stands to lose, particularly as resource value and reproductive investment likely covary with stage. Disentangling these effects is an important next step for the emerging fields of female birdsong and aggression.

We were surprised that aggression did not increase during the egg laying stage because in several respects, a female has the most to lose at this stage. The investment in the cavity remains after egg laying begins and a female evicted during egg laying must replace not only her eggs, but the resources that have accumulated over the breeding cycle: a place to deposit the eggs she is producing and a suitable cavity, which is now scarcer as the season progresses. She simultaneously risks losing her mate, her breeding status, and a developing clutch. Because both male and female house wrens commit ovicide, with successful territory replacement always involving destruction of the clutch (Belles-Isles & Picman, 1986; Pribil & Picman, 1992), we expected aggression to rise in defense of the brood. Aggression instead declined at egg laying, which suggests these resources are not simply additive and that something specific to the nest building stage drives the peak. We propose that the threat to breeding status specifically during nest building, when the two females’ broods are most likely to align, offers a particularly compelling explanation for why females were most aggressive at the nest building stage.

Lower egg laying aggression may also have physiological and ecological causes. Females may experience a decline in hormones that mediate aggression throughout egg laying. In female tree swallows, circulating testosterone levels decline between territory establishment and egg laying, likely because elevated testosterone at this stage disrupts key reproductive processes like yolk formation and incubation behavior (George et al., 2024; George & Rosvall, 2018). Additionally, aggression can become more costly once eggs are present, due to risks of injury, oxidative stress, or lost foraging time (Enquist & Leimar, 1987; Georgiev et al., 2015; Marler & Moore, 1991). Females may therefore avoid aggressive encounters that could reduce reproductive success. Similarly, work on eastern bluebirds shows that female aggression peaks prior to egg laying in order to defend nest sites and prevent egg dumping (Gowaty & Wagner, 1988). It is important to note, however, that females are not non-aggressive at the egg laying stage, as they still responded to simulated intrusions, several of which showed high aggressive responses. Because the cavity and clutch cannot be easily separated at this stage, aggression during egg laying may still defend continued access to the cavity. However, consistent with clutch-protection, Krieg and Getty (2020) found that more aggressive females during these assays at egg laying lost fewer eggs to house wren ovicide. Thus, females have good reasons to remain at least somewhat aggressive during egg laying.

### Female song as a multi-functional signal across breeding stages

Surprisingly, we did not see a significant difference of female song use during simulated intrusions across breeding stages, though it declined by the egg laying period. This is consistent with female song rates measured outside of experimental contexts in other populations, where female song is highest prelaying and increases again in early egg laying, although not as high as the initial spike (Krieg & Getty, 2016; Odom et al., 2026). This pattern may suggest that female song is not tightly linked to competition at a specific stage. Instead, it may function more generally as a flexible competitive signal used across multiple social contexts early in the breeding cycle, with reduced signaling once reproductive investment is underway. This reduction may also reflect the predation cost of signaling near an active nest. In superb fairy-wrens, female song rate is lower during incubation, and because females sing close to the nest, their song rate predicts egg and nestling predation (Kleindorfer et al., 2016). Reduced singing once a clutch is present may therefore lower the risk of revealing the nest to predators.

Substantial variation among females in singing propensity may also have contributed to the absence of clear stage differences, indicating that individual signaling strategies play an important role in shaping female song expression. When examining females that received simulated territorial intrusions at multiple stages, some individuals sang consistently across stages while others never sang at any stage. This pattern suggests the presence of “singer” and “non-singer” phenotypes within the population. Among females that did sing, song output did not differ significantly across breeding stages, further indicating that variation in female song may be driven more by individual differences than by breeding context alone. However, only a subset of females was sampled repeatedly across all stages. Future studies with larger within-individual sample sizes will be important for determining whether stage-related differences in song expression are more pronounced than what we detected.

Taken together, these findings suggest that female song in house wrens likely serves multiple functions that vary across social contexts and populations. In competitive mating contexts, for example, female song rate increases significantly upon the arrival of a second female in polygynous situations (Gleason et al., *in press*), suggesting a role in mate competition. However, research in a Michigan population points to a different primary driver: female song appears most strongly expressed during early egg laying, where it may function in preventing ovicide, defending access to the cavity, or exclusive access to the male (Krieg & Getty, 2016). Rather than contradicting our findings, this contrast highlights that the relative importance of different reproductive threats may vary across populations, shaping which contexts most strongly elicit female song. Consistent with this interpretation, ovicide rates in our study population were low compared to other populations, potentially reducing selection for signaling during egg laying. Interestingly, a recent study on natural female song rates in house wrens over a longer time course in the season found that some females showed punctuated bursts of high song output upon arrival, (Odom et al., 2026), echoing the individual variation we observed. The function of this early singing remains unclear, and future work should examine whether it reflects territory establishment, mate competition, or other early-season social interactions.

Importantly, several studies demonstrate clear fitness benefits associated with female song. In New Zealand bellbirds (*Anthornis melanura*) and superb fairy-wrens (*Malurus cyaneus)*, female song rates are correlated with reproductive success (Brunton et al., 2016; Cain et al., 2015). Similarly, female house wrens that sang more in response to intruders lost significantly fewer eggs to ovicide (Krieg & Getty, 2016). Together with our findings, this suggests that while a single resource may not directly drive female song, it likely functions as a strategic signal deployed during intrasexual competition with important fitness consequences.

Future work should therefore examine how individual variation in female song relates to reproductive outcomes, such as clutch survival, offspring condition, territory retention, or access to high-quality breeding resources.

### Female call behavior shifts across competitive contexts

Our results showed that females changed both the type and abundance of calls produced across breeding stages. Females produced chatter and harsh alarm calls most frequently during arrival and nest building, with both call types decreasing significantly by egg laying. In contrast, females produced the highest number of HI calls and sweep calls during the nest building stage. This shift in call use suggests that females adjust their vocal signaling depending on the competitive context they face during the early breeding cycle.

The high number of HI calls during nest building further corroborates our behavioral results showing that females were most aggressive during this stage. HI calls are known to be associated with aggressive interactions in house wrens (Krieg & Burnett, 2017), occurring significantly more often during playback of conspecific females compared to heterospecifics and increasing immediately prior to physical attack. The elevated production of HI calls during nest building therefore provides additional vocal evidence for the aggressive behavior pattern we observed, further supporting nest building as the stage of most intense female-female competition.

We also found that females exhibiting higher levels of aggressive behavior produced more HI calls and sweep calls overall. This relationship suggests that certain call types may function as components of aggressive signaling during competitive interactions. If calls are used in aggressive contexts, they may allow females to signal competitive ability or escalate interactions without immediately resorting to physical attack (Maynard-Smith & Harper, 2004; Searcy, 2005).

Despite evidence that some house wren calls are associated with aggressive interactions (Krieg & Burnett, 2017), the specific functions of many calls remain poorly understood. More broadly, breeding females have been shown to alter calling behavior across breeding stages, from mating through fledgling care (Amy et al., 2018; Derégnaucourt & Guyomarc’H, 2003; Halfwerk et al., 2011; Leonardi et al., 2014). These findings indicate that female call production can vary with social and reproductive context, though the functional significance of many call types remains unclear.

Furthermore, the house wren call repertoire has not been comprehensively defined, particularly in females. The call categories used in this study were based on distinct structural differences observed in spectrograms, but additional work is needed to formally characterize the repertoire and determine the functional roles of each call type. In particular, the sweep calls described here have not previously been reported in female house wren literature, though they are observable in Michigan and Pennsylvania populations as well (Krieg, pers. obs.). These calls were structurally similar to the high-pitched vocalizations males are known to produce during courtship displays, yet many females produced them during simulated competitive interactions. Interestingly, sweep calls were most strongly associated with highly aggressive females, even more so than HI calls. This raises the possibility that sweep calls may serve as an additional aggressive or competitive signal used by females during intrasexual contests. Future work should investigate the full call repertoire of house wrens and explore whether sweep calls function as an aggressive signal unique to female intrasexual competition, distinct from their structural analogs in male courtship.

### Female-female Competition

Together, these findings provide strong evidence that female-female competition plays a central role in shaping female behavior in northern house wrens, aligning with growing evidence that female–female competition is an important evolutionary driver of female elaborate traits such as aggression and song (Cain & Rosvall, 2014; Stockley & Campbell, 2013). In many systems, female competition arises because access to essential breeding resources is limited, selecting for behaviors that allow females to compete for these resources (Clutton-Brock & Huchard, 2013; Rosvall, 2011; Stockley & Campbell, 2013). In this system, aggression appears to be most strongly linked to competition over nest cavities, particularly during the nest building stage. However, in different systems, females may also compete over other resources depending on context. For example, when the operational sex ratio becomes female-biased and mates are scarce, females may compete directly for access to partners and the direct and indirect benefits they provide, particularly in species where males contribute parental care or resources (Emlen & Oring, 1977; Lewis et al., 2004; Rosvall, 2011). Similarly, when foraging resources are limited, female aggression can intensify in contexts where access to food determines reproductive success (Baird & Sloan, 2003; Ueda & Kidokoro, 2002). In other cases, females compete to defend offspring or eggs from conspecific threats (Maestripieri, 1992; Wolff & Peterson, 1998).

Ultimately, female competition over these resources has important fitness implications. Across taxa, female aggression can improve access to resources that influence breeding opportunities and reproductive success (Rosvall, 2011), consistent with broader theoretical frameworks proposing that competition for ecological and social resources can drive the evolution of traits in both sexes (Tobias et al., 2012). In northern house wrens, for example, more aggressive females produced heavier offspring and were more likely to successfully fledge young (Krieg & Getty, 2020). Together, these results suggest that the ability to compete for limiting resources can directly shape reproductive success and the evolution of female competitive traits across the breeding season.

## Conclusion

In summary, physical aggression and aggressive calls peaked during the nest building stage, identifying it as the period of most intense female-female competition in our population. This pattern is consistent with access to the nest cavity being the resource females contest most strongly, but females may also be defending breeding status and access to their built nest. In contrast, song rate did not significantly vary across stages, while the context-dependent shifts in call type reveal that females deploy different vocal signals for different competitive challenges, a flexibility that likely reflects the multifunctional nature of female communication. Notably, we noticed the presence of distinct singer and non-singer phenotypes, suggesting that individual signaling strategies may be just as important as breeding context in shaping vocal behavior, a dimension of female competition that remains poorly understood. Together, these findings challenge the assumption that female competitive behaviors are uniform and highlight that stage-specific pressures shape both the form and intensity of female signaling. Future work that tracks individual females across the full breeding cycle will be essential to unraveling how competition, communication, and individual strategy interact to shape female reproductive success.

## Data Availability

The data supporting this study are available in the Supplementary Material.

