## Supplementary Material for "A comparison of female competitive traits: Female aggression peaks at nest building but female song spans multiple contexts in a temperate songbird"

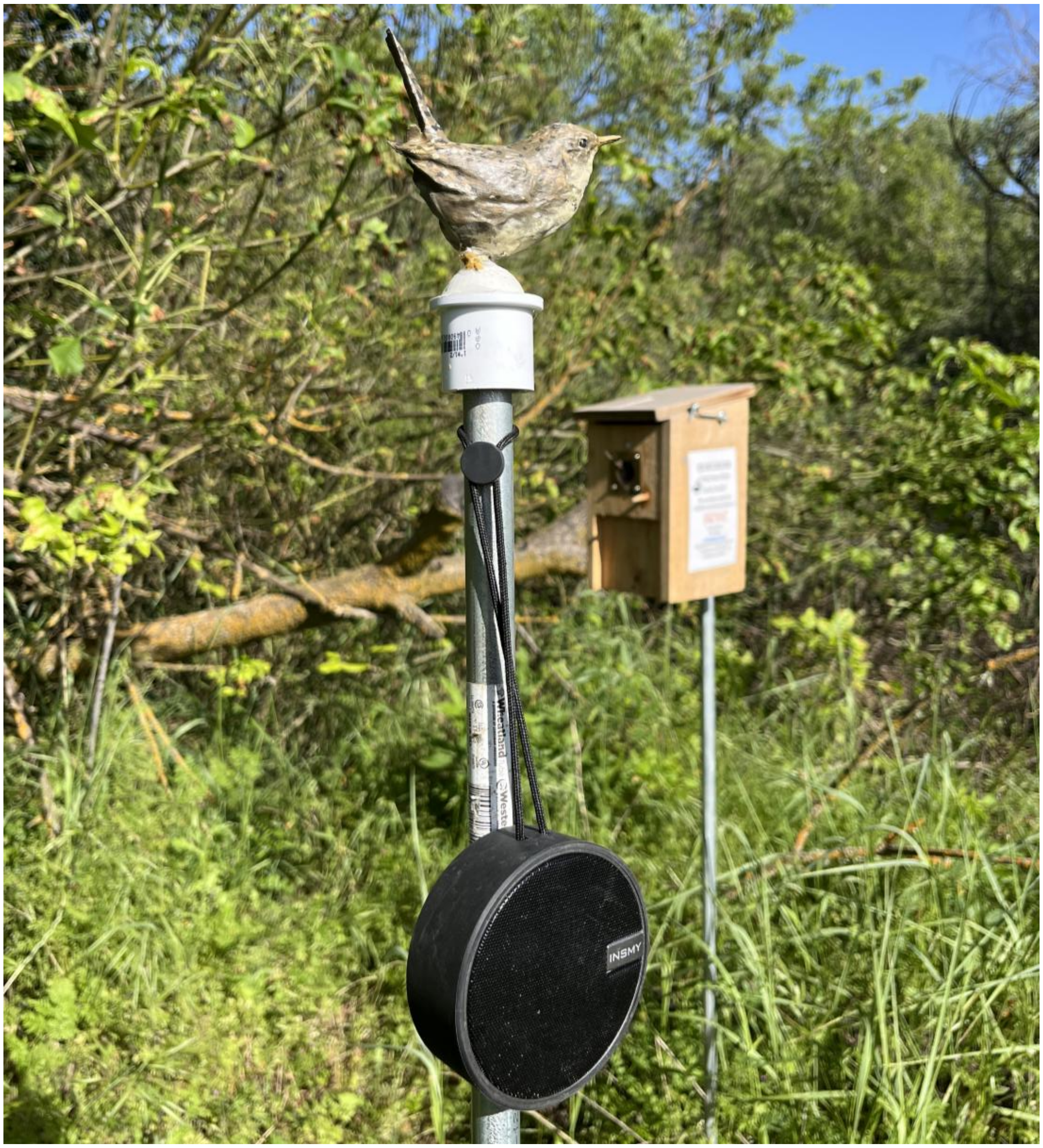


**Figure S1.** House wren model acting as visual stimuli during simulated intrusions

### Aggression

Females were classified into three ordinal aggression categories (low, medium, high) based on observed behavior during simulated territorial intrusions. PC1 scores differed significantly across aggression classes (Kruskal-Wallis: $\chi^2 = 41.47$, df = 2, $p < 0.001$), confirming that the three categories represent meaningfully distinct levels of aggressive behavior (Table S1).

**Table S1.** Aggression Metrics by Classification Level

| **Variable** | **Low** | **Medium** | **High** |
|---|---|---|---|
| Dives or attacks | 0.00 (± 0.00) | 0.00 (± 0.00) | 0.81 (± 0.08) |
| Latency to approach 5 m (s) | 600.00 (± 0.00) | 162.59 (± 42.73) | 65.97 (± 16.51) |
| Time within 1 m vs pre (s) | 0.00 (± 0.00) | 0.00 (± 0.00) | 0.92 (± 0.05) |
| PC1 score | -1.54 (± 0.00) | -0.90 (± 0.09) | 1.15 (± 0.15) |

Values are mean (± SE). Dives/attacks and time within 1 m are proportions; latency and PC1 score are continuous. PC1 derived from PCA of latency, within-1m time, and attack behavior (60.6% variance explained).

**Table S2.** PC Loading of Aggression Variables

| **Variable** | **PC1** | **PC2** | **PC3** |
|---|---|---|---|
| Latency to approach within 5 m | 0.59 | -0.53 | 0.61 |
| Approach within 1 m | 0.67 | -0.10 | -0.74 |
| Dive or attack | 0.45 | 0.84 | 0.30 |
| *Variance explained (%)* | *60.6* | *28.2* | *11.2* |

**Aggression model fit diagnostics:**

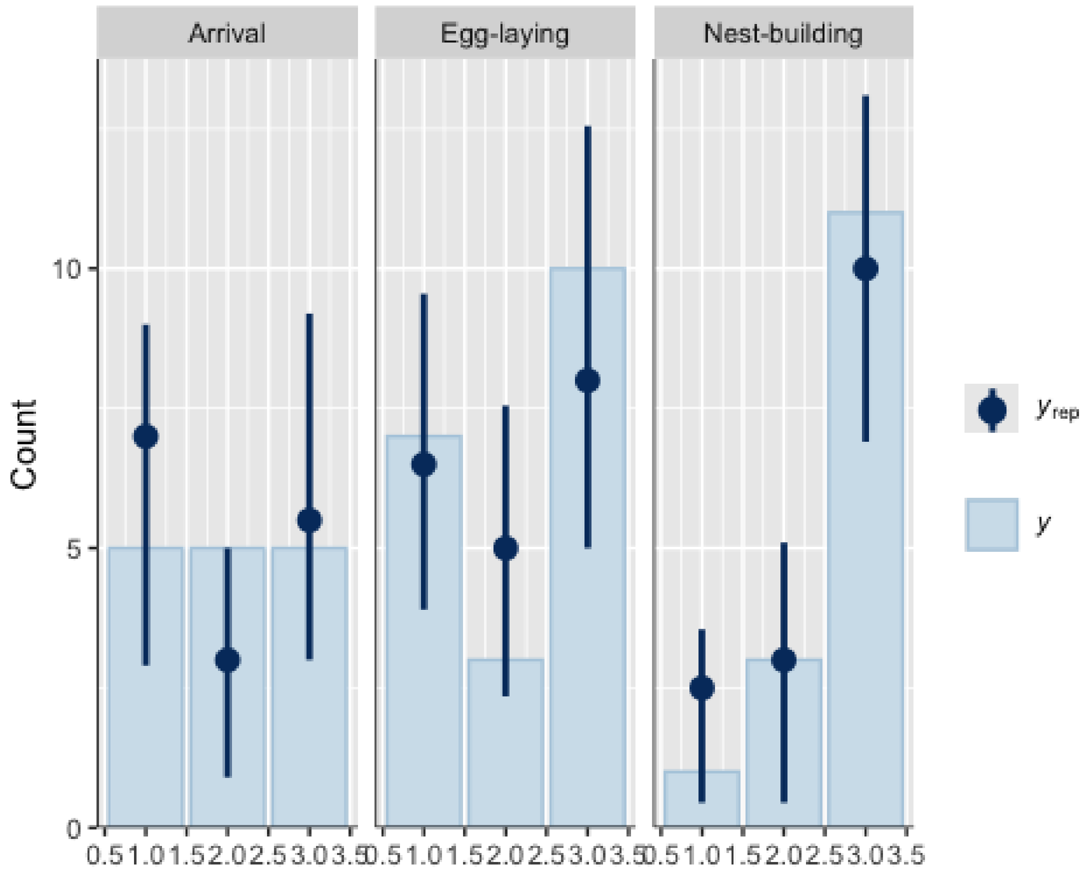


**Figure S2**. **Posterior predictive checks indicated good model fit.** The distribution of model-predicted aggression scores matched observed frequencies across all breeding stages

Posterior predictive checks indicated good model fit; the distribution of model-predicted aggression scores matched observed frequencies across breeding stages (Figure S2). We compared the base cumulative logit mixed model (breeding stage + random intercept for female identity) to seven alternative models additionally including site, exposure, and/or playback track ID as fixed or random effects. PSIS-LOO cross-validation showed no meaningful improvement in predictive performance for any alternative model (Table S3), as all ΔELPD values were within approximately one standard error of the best-fitting model. We therefore retained the base model for inference.

**Table S3**. Leave-One-Out Cross-Validation (LOO) Model Comparison for Female Aggression

| Model | ΔELPD | SE |
|---|---|---|
| **+ Track ID** | **0.0** | **--** |
| Base (stage + female ID) | -0.3 | 2.7 |
| + Site + Track ID | -0.4 | 1.4 |
| + Site | -0.6 | 3.1 |
| + Site + Trial number | -1.8 | 3.0 |
| + Trial number | -1.9 | 2.7 |
| + Trial number + Track ID | -2.0 | 0.5 |
| + All covariates | -2.2 | 1.4 |

All models include breeding stage as a fixed effect and a random intercept for female identity. ΔELPD = difference in expected log predictive density relative to the best-fitting model (+ Track ID); SE = standard error of the difference. The base model differed from the best-fitting model by only 0.3 ELPD (SE = 2.7), well within the range of sampling error, so the base model was retained for inference.

**Table S4.** Pairwise Comparisons of Female Aggression (Continuous PC1) Among Breeding Stages

| Comparison | Estimate | SE | *df* | *t* | *p* |
|---|---|---|---|---|---|
| Arrival vs. (Egg laying) | -0.02 | 0.40 | 35.3 | -0.05 | .999 |
| **Arrival vs. (Nest building)** | **-1.34** | **0.43** | **39.7** | **-3.08** | **.010** |
| **(Egg laying) vs. (Nest building)** | **-1.32** | **0.40** | **33.6** | **-3.29** | **.006** |

Estimates are differences in aggression PC1 scores from a linear mixed model with breeding stage as a fixed effect and female identity as a random effect; higher PC1 indicates greater aggression. p-values are Tukey-adjusted for multiple comparisons. Significant results are bolded.

**Female song**

**Table S5.** AIC Model Comparison for Female Song Output

| Model | df | AIC | ΔAIC |
|---|---|---|---|
| **Base + Site** | **7** | **236.92** | **0.00** |
| Base (Tweedie) | 6 | 240.21 | 3.29 |
| Base + Trial number | 7 | 241.49 | 4.57 |
| Base + Track ID | 7 | 242.21 | 5.29 |
| Base + Site + Track ID | 7 | 242.21 | 5.29 |
| Base + Track ID + Trial number | 7 | 242.21 | 5.29 |
| All covariates | 10 | 242.74 | 5.82 |
| Zero-inflated Tweedie | 7 | 242.80 | 5.88 |
| Base + Site + Trial number | 8 | 243.32 | 6.40 |
| Negative binomial | 5 | 252.20 | 15.28 |

All Tweedie models include a random intercept for female identity. ΔAIC = difference in AIC relative to the best-supported model. Models with ΔAIC > 2 have substantially less support.

**Table S6.** GLMM (Tweedie) Fixed Effects for Female Song Counts During Simulated Intrusions Across Breeding Stages

| Term | Estimate | SE | 95% CI | z | p |
|---|---|---|---|---|---|
| Intercept (Arrival) | -0.06 | 0.84 | -1.71–1.59 | -0.07 | 0.94 |
| Nest Building (vs Arrival) | -0.05 | 0.42 | -0.88–0.77 | -0.12 | 0.90 |
| **Egg Laying** (vs Arrival) | **-0.93** | **0.44** | **-1.79– -0.07** | **-2.13** | **0.033** |
| **Site** | **1.78** | **0.81** | **0.19–3.37** | **2.19** | **0.029** |

Estimates are on the log scale. Female identity was included as a random effect and field site as a fixed effect to account for location differences. Bolded rows indicate $p < 0.05$.

**Calls**

**Table S7.** SS Loadings and Variance Explained by Retained Principal Components from the Promax-Rotated PCA

| PC | SS Loadings | Variance explained (%) | Cumulative variance (%) | Primary contributors (\|loading\| ≥ 0.40) |
|---|---|---|---|---|
| PC1 | 1.810 | 36.2 | 36.2 | Sweeps, HI, Harsh |
| PC2 | 1.196 | 23.9 | 60.1 | Chatter, Harsh |
| PC3 | 1.117 | 22.3 | 82.5 | Other |

SS Loadings = sum of squared loadings per component after Promax rotation. Variance explained is relative to total variance across all variables.

**Table S8.** Promax-Rotated Loadings for PC1-3

| Vocalization | PC1 | PC2 | PC3 | Primary PC | Direction |
|---|---|---|---|---|---|
| HI call | 0.82 | 0.00 | -0.17 | PC1 | + |
| Sweep | 0.93 | -0.21 | 0.30 | PC1 | + |
| Harsh | 0.54 | 0.41 | -0.18 | PC1 | + |
| Chatter | -0.12 | 1.02 | 0.16 | PC2 | + |
| Other | 0.07 | 0.15 | 0.99 | PC3 | + |

Cells Loading >= 0.40 Are Highlighted.

**Relationship between aggression and female vocalizations**

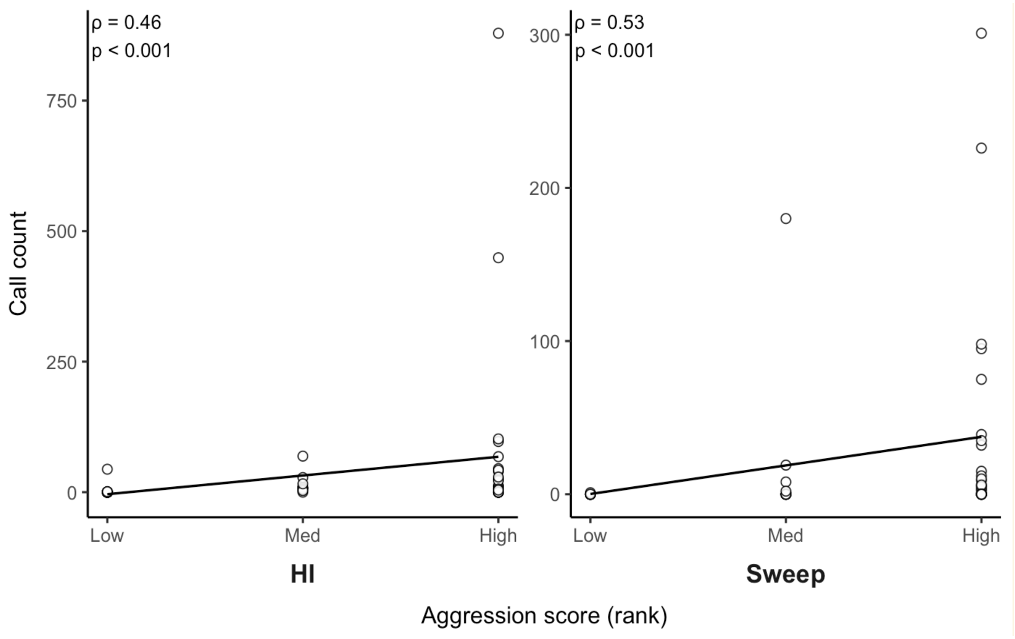


**Figure S3. The Relationship Between Aggression Scores and Call Production During Simulated Intrusions.** Call counts of a) HI calls and b) Sweep calls plotted against original aggression scores (low, medium, high). Points represent individual trials; solid lines indicate trends. Associations were assessed using Spearman's rank correlation with corresponding correlation coefficient and p-values shown.
